# The serotonergic psychedelic DOI impairs deviance detection in the auditory cortex

**DOI:** 10.1101/2024.09.06.611733

**Authors:** Max Horrocks, Jennifer L. Mohn, Santiago Jaramillo

## Abstract

Psychedelics are known to induce profound perceptual distortions, yet the neural mechanisms underlying these effects, particularly within the auditory system, remain poorly understood. In this study, we investigated the effects of the psychedelic compound 2,5-Dimethoxy-4-iodoamphetamine (DOI), a serotonin 2A receptor agonist, on the activity of neurons in the auditory cortex of awake mice. We examined whether DOI administration alters sound-frequency tuning, variability in neural responses, and deviance detection (a neural process reflecting the balance between top-down and bottom-up processing). Our results show that while DOI does not alter the frequency selectivity of auditory cortical neurons in a consistent manner, it increases trial-by-trial variability in responses and consistently diminishes the neural distinction between expected (standard) and unexpected (oddball) stimuli. This reduction in deviance detection was primarily driven by a decrease in the response to oddball sounds, suggesting that DOI dampens the auditory cortex’s sensitivity to unexpected events. These findings provide insights into how psychedelics disrupt sensory processing and shed light on the neural mechanisms underlying the altered perception of auditory stimuli observed in the psychedelic state.

**New & Noteworthy:** The neural basis of perceptual distortions induced by psychedelics remains poorly understood. This study demonstrates that the serotonergic psychedelic DOI increases neural response variability and impairs deviance detection in the auditory cortex by reducing sensitivity to unexpected sounds. These findings provide new insights into how psychedelics disrupt sensory processing and alter the balance between bottom-up and top-down neural signaling, contributing to our understanding of altered perception in the psychedelic state.

## Introduction

Psychedelics, a subclass of hallucinogenic psychoactive compounds, have long fascinated humanity by offering profound and often transformative experiences that alter perception, cognition, and mood (1, 2). Additionally, there has been a recent resurgence in the use of psychedelics as therapeutic agents for the treatment of conditions such as depression, anxiety, and trauma-related disorders (3, 4). These factors underscore the importance of understanding the neural mechanisms underlying the perceptual changes induced by these substances.

Serotonergic psychedelics primarily exert their effects through the serotonin 2A (5-HT2A) receptor (5), leading to various perceptual distortions across sensory systems. For classical psychedelics, such as LSD and psilocybin, perceptual distortions are dominated by visual phenomena, such as vividly colored, constantly changing geometric figures that can evolve into complex images and scenes at medium to high doses. True hallucinations, defined as perceptions that feel real despite the absence of external stimuli, are less common than perceptual distortions but can occur at higher doses. While effects in auditory perception during the psychedelic state are generally reported to be less dominant than their visual counterparts, psychedelics can produce strong distortions in the passing of time and in the perception of sounds (6, 7, 8), suggesting direct effects onto the auditory system.

Studies using human subjects provide invaluable information about the psychological effects of different classes of psychoactive compounds, yet a detailed understanding of the neural mechanisms underlying perceptual distortions requires the level of experimental access that animal models provide. Studies in animals have provided key insights into the molecular mechanisms of action of psychedelics (9), however, our understanding of how psychedelics influence neuronal activity in the auditory system to produce these perceptual distortions remains limited. Several processes have been proposed for the distortions observed during the psychedelic state, including disruption of thalamocortical connectivity, altered cortical network dynamics, and impacts on the default-mode network (10, 11, 12). A prevailing theory, compatible with the aforementioned mechanisms, suggests that psychedelics induce perceptual changes by altering the balance between top-down and bottom-up processing in the brain. For instance, researchers have proposed that psychedelics result in a relax of top-down control, leading to enhanced bottom-up sensory processing and novel perceptual experiences (13).

Here, we use systemic administration of the psychedelic 2,5-Dimethoxy-4-iodoamphetamine (DOI) to evaluate the effects of the psychedelic state on the representation of sounds by auditory cortical neurons. We first test whether DOI affects the overall activity and/or selectivity to sound frequency of auditory cortical neurons. We then characterize the effects of DOI on a neural phenomenon that depends on the balance between bottom-up and top-down processes, namely that of deviance detection (14). Our findings reveal that while DOI administration results in no net effect in frequency-tuning across the population of auditory cortical neurons, it consistently diminishes the response difference between expected and unexpected stimuli in most of these neurons.

## Materials and Methods

### Animals

Four adult C57BL/6J mice (JAX 000664, RRID:IMSR JAX:000664), two male and two female, were used for electrophysiological recordings. At the time of the recordings, three mice were 20 weeks old and one mouse was 28 weeks old. Two additional male mice of the same strain (19 weeks old) were used to evaluate head-twitch responses under DOI. All procedures followed the National Institutes of Health animal care standards and were approved by the University of Oregon Animal Care and Use Committee.

### Sound presentation

Experiments were performed inside a single-walled sound-isolation box (IAC Acoustics, North Aurora, IL). Auditory stimuli were presented in an open-field configuration from a speaker (MF1, Tucker-Davis Technologies, Alachua, FL) contralateral to the side of electrophysiological recordings. Speakers were calibrated using an ultrasonic microphone (ANL-940-1, Med Associates, Inc., Fairfax, VT) to obtain the desired sound intensity level for frequencies between 1 kHz and 40 kHz. Stimuli were generated using Python software developed in-house (https://taskontrol.readthedocs.io/). During sound presentation, mice were awake and head-fixed on top of a freely-moving wheel, leaving them free to move their limbs while their heads remained stationary.

Three different ensembles of stimuli were used in our experiments: one for evaluating frequency tuning, and two different ensembles for evaluating deviance detection. To evaluate frequency tuning, we used pure-tone pips (100 ms duration) at 16 frequencies logarithmically-spaced between 2 kHz and 40 kHz at 70 dB SPL (with a 2 ms amplitude ramp up and ramp down) presented in random order. A session consisted of at least 20 repetitions per frequency with interstimulus intervals randomized in the range 1–1.4 seconds. For the first type of oddball paradigm, we used chords composed of 12 simultaneous pure tones (50 dB SPL each) logarithmically spaced in the range *f /*1.2 to *f ×* 1.2, for a given center frequency *f*. Two center frequencies, 8 kHz and 13 kHz were used. The sequence consisted of sounds 50 ms long spaced by 500 ms of silence, where one frequency (the standard) was repeated nine to eleven times followed by a single presentation of the other frequency (the oddball) followed by another sequence of standards and so on. After the presentation of at least 50 oddball stimuli, the sequence was repeated, but exchanging which stimulus served as the standard. For the second type of oddball paradigm, we used frequency modulated (FM) sounds consisting of either an up-sweep between 8 and 13 kHz or a down-sweep between 13 and 8 kHz, at an intensity of 70 dB SPL. Sounds in the sequence were 100 ms long spaced by 500 ms silence, and followed a repetition rate similar to the one for chords (nine to eleven standards for each oddball). After the presentation of at least 50 oddball stimuli, the sequence was repeated, but exchanging which stimulus served as the standard.

### Surgical procedure

Animals were anesthetized with isoflurane administered via a nose-cone while positioned on a stereotaxic apparatus. Mice were surgically implanted with a head-bar and a Neuropixels 1.0 probe in the auditory cortex (two mice on the left hemisphere and two mice on the right hemisphere), using a design based on (15). The craniotomy was centered 2.65 mm posterior to Bregma and 4.4 mm lateral to the midline. The probe was lowered 3 mm from the brain surface, and it spanned multiple subregions of the auditory cortex. All animals were monitored after surgery and recovered fully before electrophysiological experiments.

### Electrophysiological recordings and administration of DOI

Electrical signals were collected using Neuropixels 1.0 probes (IMEC) via an NI PXIe-8381 acquisition module and the Open Ephys GUI software (www.open-ephys.org, RRID:SCR 021624). During the experiment, mice were head-fixed and allowed to run on top of a wheel. We performed electro-physiological recordings on five different days for each mouse. There was no formal power analysis performed to determine the sample size, but we estimated that our number of animals and sessions would yield about 100 neurons per condition given the strict criteria for inclusion of cells in the analysis described below. Each recording day consisted of multiple sessions before and after injections of reagents and while presenting the stimulus ensembles described above. We began with electro-physiological recordings before administering any reagents, during which we presented three stimulus ensembles to evaluate frequency tuning and deviance detection. We then injected a volume of 5 ml/kg of sterile saline (0.9% concentration) subcutaneously and waited 15 minutes before another set of recordings during the presentation of all three stimulus ensembles. Finally, we injected 2,5-Dimethoxy-4-iodoamphetamine, (±)-DOI hydrochloride (Sigma-Aldrich, D101) subcutaneously at a concentration of 10 mg/kg (diluted in sterile saline), and waited another 15 minutes before the last set of recordings during the presentation of all three stimulus ensembles. The concentration of DOI used here for subcutaneous delivery matches that used in previous studies of the effects of DOI on sensory processing (16). This concentration is intentionally higher that doses used with intraperitoneal injections of DOI (17, 18) to account for differences in pharmacokinetics between these methods of administration (19). There was no blinding regarding the reagent used in each part of the experiment.

### Estimation of recording location

Before inserting the Neuropixels probes intro the brain, the probes were coated with a fluorescent dye (DiI: Cat# V22885, Thermo-Fisher Scientific) to allow for identification of the recording locations post-mortem (Fig. 1A). At the conclusion of the experiments, animals were euthanized with euthasol and perfused through the heart with 4% paraformaldehyde. Brains were extracted and left in 4% paraformaldehyde for at least 24 hours before slicing. Brain slices (thickness 100 *µ*m) were prepared under phosphate-buffered saline using a vibratome (Leica VT1000 S) and imaged using a fluorescence microscope (Axio Imager 2, Carl Zeiss) with a 2.5x objective. To determine the location of our recording sites, we manually registered each brain slice to the corresponding coronal section in Allen Mouse Brain Common Coordinate Framework (CCF) (20), and estimated the position of the electrode that showed the strongest signal for each neuron.

**Figure 1:**
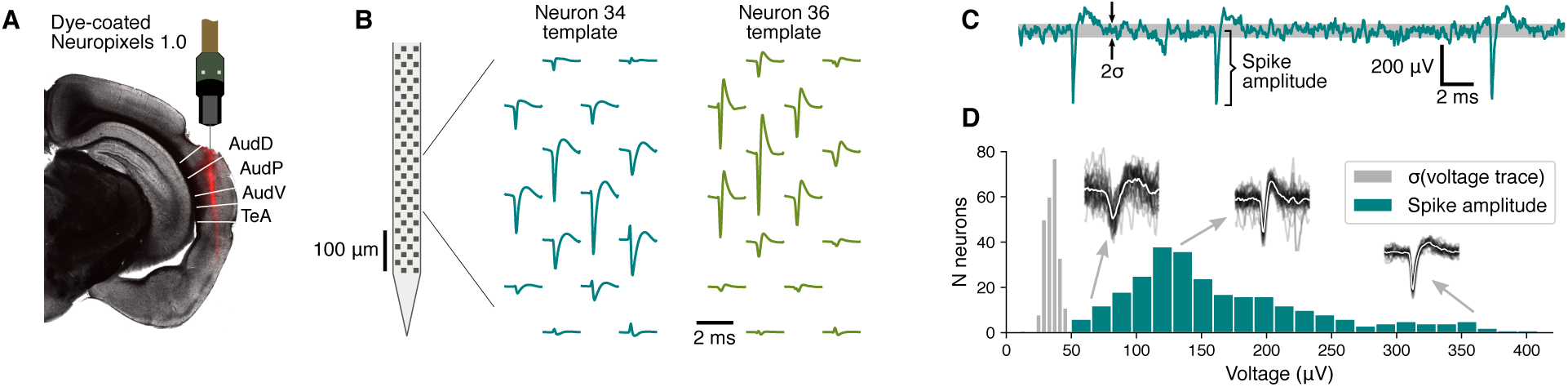
Neuropixels recordings from the auditory cortex. **(A)** Coronal slice of mouse brain showing the red track left by a dye-coated Neuropixels probe used for recording neural activity. The penetration spanned multiple fields of the auditory cortex (Dorsal, Primary, and Ventral fields) and the temporal association cortex (TeA). **(B)** Two examples of templates estimated by the spike sorting software package Kilosort. Both neurons showed their largest signals around the same recording electrodes in the probes, but had very different spike shapes. **(C)** Voltage trace for one recording electrode showing large spikes. The baseline noise level is illustrated by a gray band at one standard deviation (*σ*) above and below the mean of the signal. **(D)** Distribution of average spike amplitudes of neurons included in the study (measured on the channel with largest signal for each neuron), compared to the distribution of noise levels observed. Insets show examples of spikes from three different neurons of different quality, not at scale (black: 30 sample traces, white: average trace).

### Estimation of locomotion state

The locomotion state of the animal (running *vs.* not-running) was estimated from video recordings during each session. Videos were collected with a USB infrared camera at 30 frames per second and processed using the software package FaceMap v0.2 (21) (RRID:SCR 021513) to extract the average change in pixel intensity in the area containing the animal’s paws. The trace of pixel intensity change was smoothed using a 10-frame square window, and the value at stimulus onset was thresholded to determine whether the animal was in a running or not-running state.

### Analysis of neural data

The spiking activity of single units was isolated using the spike sorting package Kilosort v2.5 (22) (RRID:SCR 016422) and manually curated using the software Phy (23) (Fig. 1). The resulting data were analyzed using in-house software (https://github.com/sjara/jaratoolbox) developed in Python (www.python.org, RRID:SCR 008394). To ensure that differences in neural responses between conditions were not merely due to the passage of time in the rig, we focused all analysis on neurons for which the cell’s firing, spontaneous or evoked, did not change more than 30% between the pre-injection and the saline conditions. Responsiveness to sounds was evaluated by calculating whether there was a statistically significant difference between evoked firing (throughout the duration of each sound) and baseline activity (−200 ms to 0 ms from sound onset), for any of the sounds presented, using a Wilcoxon signed-rank test with Bonferroni correction for multiple-comparisons. Evoked firing was evaluated in the period from 15 ms after stimulus onset to 15 ms after stimulus offset to account for the signal transmission delay from the ear to the auditory cortex.

To evaluate the pure-tone frequency selectivity of a neuron, a Gaussian function was fit to the spike counts in response to each frequency (using a logarithmic space for frequency). The best frequency (BF) of a neuron was defined as the peak of this Gaussian function. The width of tuning was defined as the full width at half maximum of the normalized Gaussian, estimated as 2.355*σ*, where *σ* is the standard deviation of the Gaussian. The maximum change in firing (Max Δ firing) was estimated as the difference between the Gaussian peak and the baseline firing rate.

To estimate differences between the responses to oddball and standard sounds, we defined an Oddball Enhancement Index, OEI = (*O−S*)*/*(*O* + *S*), where *O* is the average response to the oddball and *S* is the average response to the standard. To quantify these effects of a specific reagent on the responses to either oddball or standard sounds, we defined a modulation index MI = (DOI *−* Sal)*/*(DOI + Sal), where DOI and Sal are the average responses under DOI and Saline, respectively.

To estimate the trial-by-trial variability in neural responses, we calculated the Fano Factor for the responses to each pure tone for each neuron: FF = *σ* ^2^*/µ*, where *σ* ^2^ is the variance of the evoked firing rate across trials and *µ* is the mean. We then calculated the average FF across stimuli for each neuron before comparing these values between saline and DOI conditions. To obtain reliable values of the FF when analyzing running and stationary trials separately, we included in the FF average only those stimuli with at least four trials in each running condition.

### Statistical analysis

Non-parametric tests were used throughout the study to evaluate differences in neural responses under different conditions, as indicated in the main text. We used a Wilcoxon signed-rank test to determine whether neurons were responsive to a particular sound, treating the spike counts during baseline and during the stimulus as matched samples for each trial. We used a Wilcoxon signed-rank test to determine differences between Saline and DOI conditions, treating the average response of a neuron under each condition as matched samples. To compare the effect of DOI between neurons recorded during the first sessions and those recorded during the last sessions, we used the Mann-Whitney U rank test.

## Results

### DOI decreased spontaneous and evoked firing in AC

To evaluate the effects of the hallucinogen 2,5-Dimethoxy-4-iodoamphetamine (DOI) on the representation of sounds by auditory cortical neurons, we recorded neural activity from awake head-fixed mice using Neuropixels probes while animals were presented with auditory stimuli before and after subcutaneous injection of either saline (as control) or 10 mg/kg of DOI. Each experimental session consisted of three periods: one before saline injection, one after saline injection but before DOI injection, and one after DOI injection (Fig. 2A). During each period, we presented pure tones and frequency-modulated sounds to evaluate the overall responsiveness, frequency tuning, and deviance detection for each recorded neuron. We recorded five experimental sessions from each of the four mice. To confirm the effectiveness of the dose used in our experiments, we measured the rate of head-twitch responses (HTR) after injection of DOI (18, 24) in an additional group of two mice, freely moving in their cage. Twenty minutes after injection of DOI, these mice showed a rate of 2 HTR/min and 1.8 HTR/min, compared to 0.2 HTR/min for either mouse mice after saline injection.

**Figure 2:**
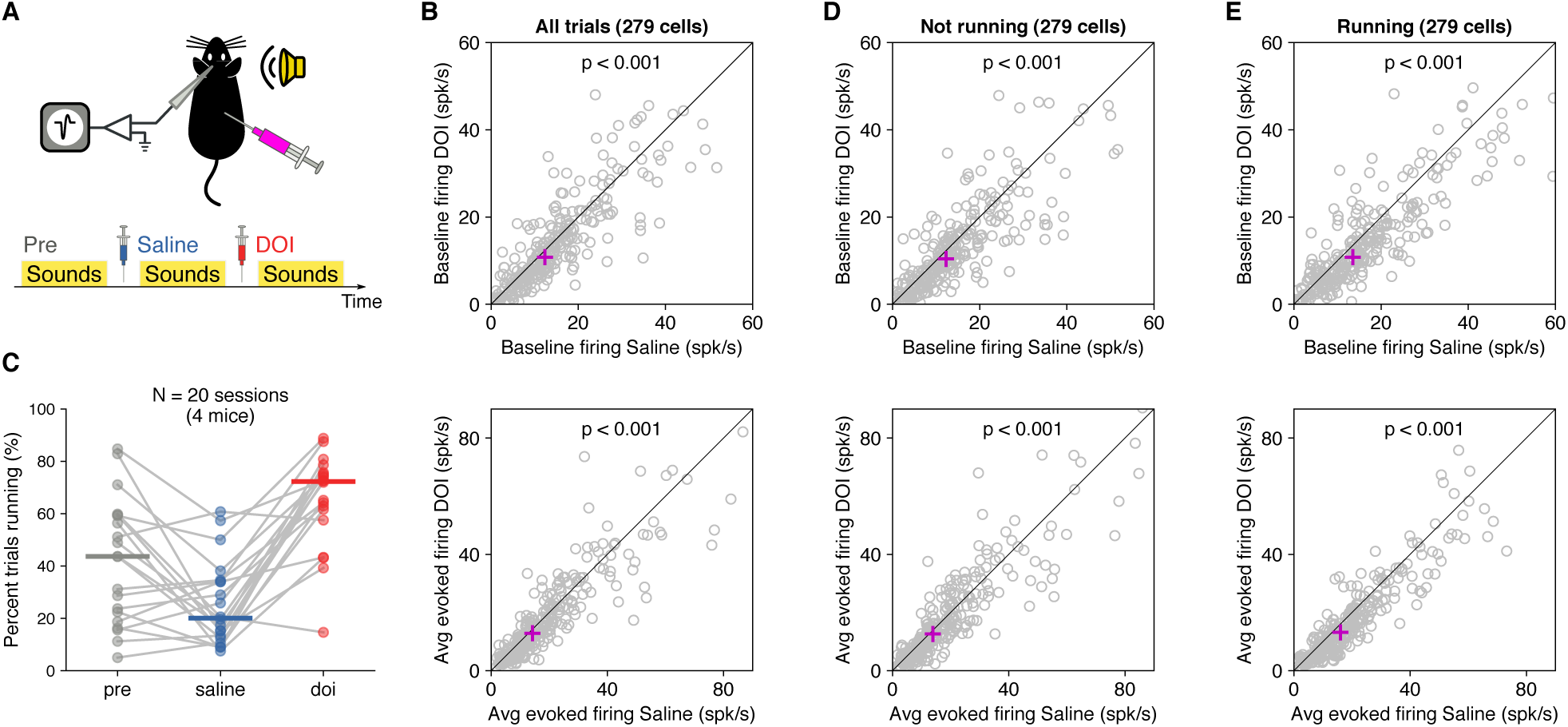
DOI decreased spontaneous and sound-evoked firing in the auditory cortex. **(A)** We recorded neural activity from the auditory cortex of awake, head-fixed mice during the presentation of auditory stimuli. Each experimental session consisted of three periods of sound presentation: one before saline injection, one after saline injection, and one after DOI injection. **(B)** Average spontaneous (top) and sound-evoked (bottom) neural activity decreased after DOI. Each dot represents one neuron. The magenta cross indicates the median value across neurons. **(C)** The percentage of trials animals spent running increased after DOI injection. Each dot represents one period for one animal, with periods from the same session connected by lines. **(D)** Same as (B) but including only trials where the animal was stationary. The decrease in spontaneous and sound-evoked neural activity remains apparent. **(E)** Same as (B) but including only trials where the animal was running. The decrease in spontaneous and sound-evoked neural activity remains apparent.

We first tested the effect of DOI on the overall firing rate of each recorded neuron. To ensure any results would not be simply due to the passage of time during the experimental session, we focused our analysis on neurons that passed two criteria: (1) the cell’s firing, spontaneous or evoked, did not change by more than 30% between the pre-injection and the saline conditions, and (2) the cell was responsive to pure tones during all three periods, with an evoked firing rate above five spikes per second. These strict criteria ensured that only cells that were stable during our recordings were included in the analysis. From the 1,590 isolated single neurons during our recordings, 279 passed these criteria. Approximate estimates of the location of these sound-responsive neurons, based on the Allen Mouse Brain Common Coordinate Framework (CCF) (20), yielded 84% of them in the main fields of the auditor cortex (42% in the primary field, 34% in the dorsal field, and 8% in the ventral field), 9% in the temporal association area, and 7% in other neighboring cortical areas. We found that, on average, the population of neurons showed a decrease in both baseline spontaneous firing (Fig. 2B top) and sound-evoked firing (Fig. 2B bottom) after DOI injection compared to the saline control period (*p <* 10*^−^*^5^ for both spontaneous and sound-evoked conditions, Wilcoxon signed-rank test).

Because psychedelics like DOI can cause changes in locomotion (25), and locomotion can alter sound-evoked responses in auditory cortical neurons (26), we measured the effect of DOI on running behavior in the head-fixed animals. We found that after DOI injection, animals were more likely to run on the wheel, with median percentages of trials (across sessions) of 44% before injections, 20% after saline, and 72% after DOI. However, we found that the effects of DOI on auditory cortical neurons could not be simply explained by the increase in running during DOI trials, since similar results were obtained from analyses of both stationary (Fig. 2D) and running (Fig. 2E) trials (*p <* 10*^−^*^4^ for all conditions tested, Wilcoxon signed-rank test).

### DOI increased the trial-to-trial variability in neural responses

Sensory neurons typically exhibit some variability in their responses to repeated presentations of the same stimulus. To assess whether DOI influenced this inherent trial-to-trial variability, we measured the Fano Factor (FF) for responses to each pure tone presented before and after DOI administration. For each cell, we estimated an average FF across tone frequencies and compared this measure of response variability between the saline and DOI conditions. We found that the median FF increased from 0.81 under saline to 0.87 under DOI (*p <* 10*^−^*^6^, Wilcoxon signed-rank test), with 65.2% of neurons exhibiting a higher FF under DOI compared to saline.

We next tested whether this increase in response variability was present during both running and stationary trials. Because animals occasionally had very few trials in a specific running condition during some sessions, we limited our analysis to stimuli with at least four trials in both running conditions. We found that the difference in response variability was clearly apparent during running trials, with the median FF increasing from 0.66 under saline to 0.79 under DOI (p*<* 10^6^, Wilcoxon signed-rank test) and 65.9% of cells showing larger values under DOI compared to saline during running trials. In contrast, we found little difference in response variability during trials when animals were stationary, with median FF of 0.72 for both saline and DOI (*p* = 0.54, Wilcoxon signed-rank test) and about 47.7% of cells showing larger values under DOI compared to saline during not-running trials. These results suggest that while DOI increases the overall trial-by-trial variability of responses, these effects may vary depending on the state of the animal.

### DOI did not change the frequency tuning of AC neurons in a consistent manner

We next tested whether DOI changed the frequency tuning of neurons in the auditory cortex. For each neuron, we estimated the responses to randomly presented pure tones (100 ms) of frequencies ranging from 2 kHz to 40 kHz. Consistent with the overall changes in firing quantified above, we found neurons that showed a decrease in evoked response to pure tones after DOI (Fig. 3A), neurons that had no change in firing across frequencies (Fig. 3B), and neurons that showed an increase in evoked responses after DOI (Fig. 3C). However, overall changes in frequency tuning curves seemed minimal. To quantify potential changes in frequency tuning, we fit a Gaussian curve to the evoked responses of each neuron and measured 3 parameters of this curve: (1) the best frequency (BF), (2) the width of tuning, and (3) the maximum change in firing evoked by the presentation of pure tones (Max Δ firing) (Fig. 3D). For this analysis, we focused on neurons that showed a positive responses to pure tones, and had a reasonably good Gaussian fit (*R*^2^ *>* 0.05). A total of 197 neurons passed these criteria.

**Figure 3:**
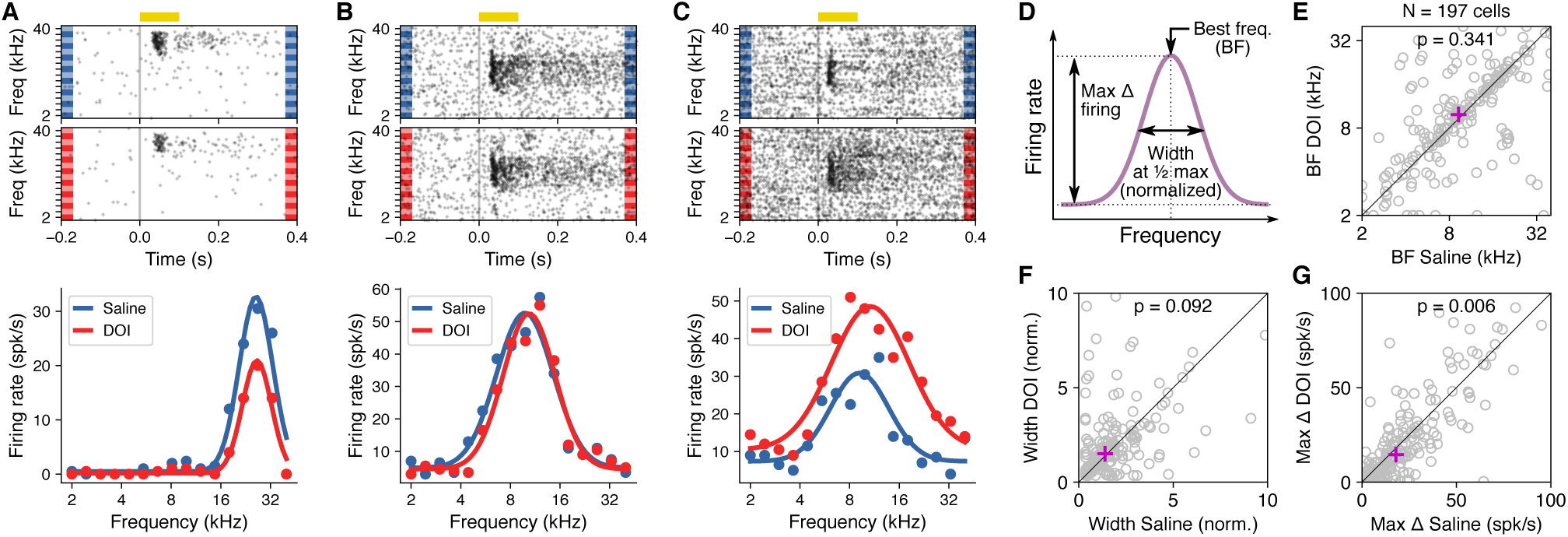
DOI did not change the frequency tuning of AC neurons in a consistent manner. **(A)** Frequency tuning for an example neuron under saline or DOI. Sound-evoked responses decrease under DOI, without a clear change in frequency tuning. **(B)** Example neuron showing no change in responsiveness or frequency tuning under DOI. **(C)** Example neuron showing an increase in sound-evoked responses under DOI, and minimal change in frequency tuning. **(D)** Illustration of the parameters of frequency tuning quantified from a Gaussian fit for each neuron: best frequency (BF), width of frequency tuning at half maximum normalized by Gaussian amplitude (Width), and maximum change in firing (Max Δ firing). **(E)** Best frequency for each neuron under DOI *vs.* saline. Magenta cross shows the median. There is no consistent change in best frequency. **(F)** Tuning width for each neuron under DOI *vs.* saline. There is no consistent change in tuning width. **(G)** Maximum change in firing evoked by sounds for each neuron under DOI *vs.* saline. There is a consistent decrease in the maximum firing after DOI.

We found that while the estimated best frequency changed for several neurons after DOI, these effects did not appear to follow a consistent direction, resulting in both increases and decreases in BF across the population (Fig. 3E) with no change in the median across neurons (*p* = 0.341, Wilcoxon signed-rank test). Similarly, we found no difference in the median values of frequency-tuning width (Fig. 3F) between saline and DOI conditions (*p* = 0.092, Wilcoxon signed-rank test). Moreover, we found no correlation between the BF or tuning width for a neuron during saline and the change in evoked response magnitude after DOI (*r* = 0.11*, p* = 0.13 for BF; *r* = 0.037*, p* = 0.61 for tuning width, Pearson correlation). The only consistent change we observed when quantifying responses to pure tones was a decrease in the magnitude of the evoked responses after DOI, measured as the height of the Gaussian fit (Fig. 3G). The median change across neurons was statistically significant (*p* = 0.006, Wilcoxon signed-rank test), aligned with the reduction in sound-evoked firing rates described earlier.

### DOI reduced sensitivity to oddball sounds

A robust and widespread phenomenon observed in sensory systems is an increased neural response to an infrequently occurring stimulus compared to the response to the same stimulus when it appears repeatedly (27, 14). Here, we tested whether DOI affects this deviance detection phenomenon by evaluating the difference in responses to a sound appearing frequently in a sequence *vs.* the same sound occurring rarely in another sequence (Fig. 4).

**Figure 4:**
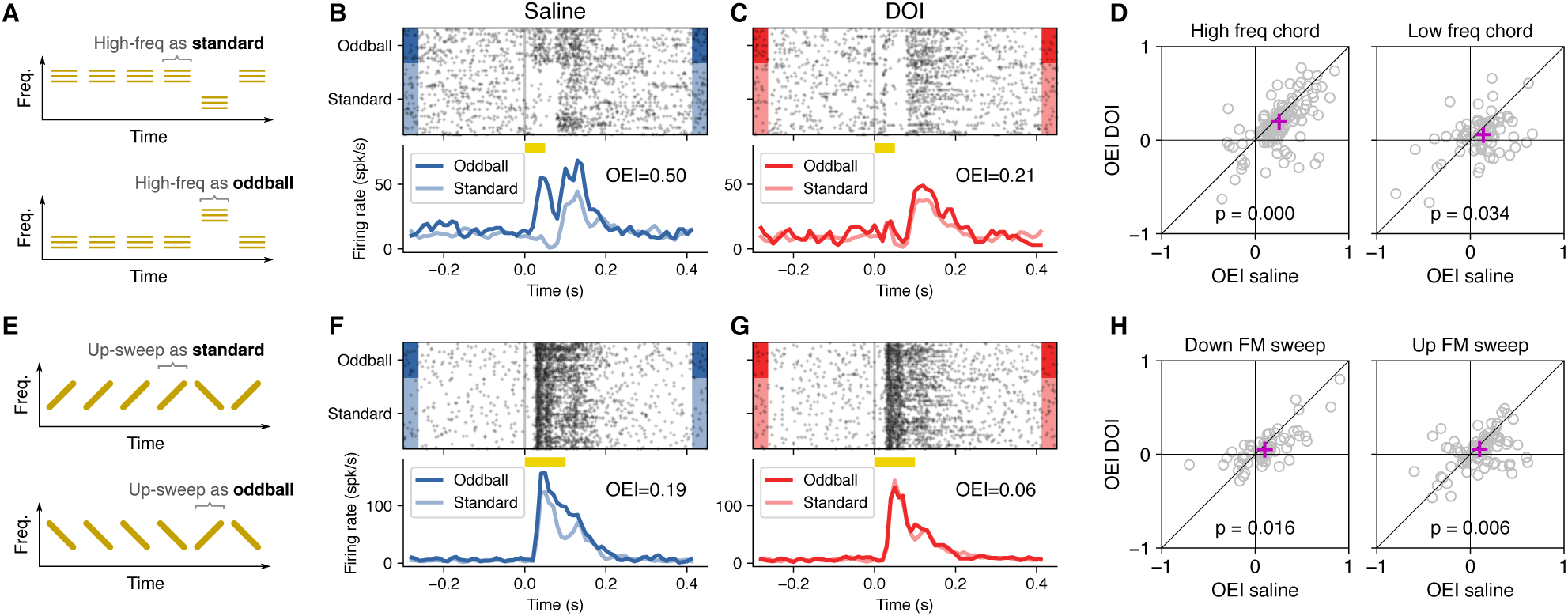
DOI reduced the difference between responses to oddballs and to standards. **(A)** Sequences of chords. The high-frequency chord appeared either as the standard (top) or as the oddball (bottom). **(B)** Responses to the high-frequency chord from an example AC neuron under the saline condition. The response to the sound was larger when it appeared as oddball. **(C)** Responses to the high-frequency chord from the example neuron in (B) after DOI injection. The difference between responses to oddball and standard was greatly reduced. **(D)** The oddball enhancement index (OEI) was smaller after DOI injection compared to the saline condition for both high- and low-frequency chords. Each dot represents one neuron. The magenta cross indicates the median value across neurons. **(E)** Sequences of frequency modulated (FM) sounds. The up-sweep sound appeared either as the standard (top) or as the oddball (bottom). **(F)** Responses to the up-sweep FM sound from an example neuron under the saline condition. The response to the sound was larger when it appeared as oddball. **(G)** Responses to the up-sweep FM sound from the example neuron in (F) after DOI injection. The difference between responses to oddball and standard was greatly reduced. **(H)** The oddball enhancement index (OEI) was smaller after DOI injection compared to the saline condition for both down- and up-sweep sounds. Each dot represents one neuron. The magenta cross indicates the median value across neurons.

We first used narrowband chords (50 ms duration) centered at two different frequencies (8 and 13 kHz) to create a sequence of sounds where one of the chords (the standard) appears repeatedly, while the other chord (the oddball) appears once every 9 to 11 presentations. In one recording session, the high-frequency chord was presented as the standard, while in a subsequent session, it was presented as the oddball (Fig. 4A). As expected for the saline control condition, we found that neural responses to a sound appearing as oddball were on average stronger than responses to the same sound appearing as standard (Fig. 4B). After DOI injection, however, this difference in responses between oddball and standard was greatly reduced (Fig. 4C).

To quantify this effect across the population of recorded neurons, we calculated for each neuron an Oddball Enhancement Index (OEI) defined as (*O−S*)*/*(*O* + *S*), where *O* is the average response to the oddball and *S* is the average response to the standard. For this analysis, we included only cells that met two criteria: (1) they were responsive to the evaluated stimulus, and (2) their OEI varied by less than 30% between the pre-injection and the saline conditions. We found that the median OEI across the population of neurons was significantly lower under DOI compared to saline (Fig. 4D). This effect was observed for both the high-frequency chord (*N* = 128 cells, *p* < 10^−5^, Wilcoxon signed-rank test) and the low-frequency chord (*N* = 79 cells, *p* = 0.034, Wilcoxon signed-rank test).

Previous studies have shown that the auditory cortex of rodents plays a larger role in processing frequency-changing sounds compared to sounds of constant frequency (28, 29, 30). With this in mind, we next aimed to test the effects of DOI on deviance detection phenomena using stimuli that are more likely to engage high-level processing in the auditory cortex. To this end, we created sequences of standards and oddballs with two frequency-modulated (FM) sounds, one up-sweep and one down-sweep, where the frequency range (8–13 kHz) and duration (100 ms) were identical. Similar to the results obtained with sequences of chords, we found a reliably positive OEI under saline (Fig. 4F) that decreased under DOI (Fig. 4G). Consistent with the results for chords, the median OEI across the population of neurons was diminished for down-sweeps (*N* = 65 cells, *p* = 0.016, Wilcoxon signed-rank test) and for up-sweeps (*N* = 95 cells, *p* = 0.006, Wilcoxon signed-rank test) (Fig. 4H).

The difference in OEI observed between saline and DOI conditions could result from either a reduction in responses to oddballs or an enhancement of responses to standards. Fig. 5A shows the data for the same example cell presented in Fig. 4F and Fig. 4G, but with responses to standard stimuli (top) and oddball stimuli (bottom) shown separately. This example suggests that the observed changes in OEI after DOI were primarily due to a decrease in responses to the oddball, rather than an increase in responses to the standard. To quantify these effects across the population of neurons, we calculated a modulation index, MI = (DOI *−* Sal)*/*(DOI+Sal), separately for standards and for oddballs. Across the population, we found that the median MI was negative for both oddball and standards (consistent with the overall reduction in responsiveness described above), but that the change in response to the oddball was larger than the change in responses to the standard. This pattern held true for chords (p*<* 10*^−^*^5^ for high-freq and *p* = 0.039 for low-freq, Wilcoxon signed-rank test)(Fig. 5B) and for FM sounds (p=0.015 for down-sweeps and *p* = 0.006 for up-sweeps, Wilcoxon signed-rank test)(Fig. 5C).

**Figure 5:**
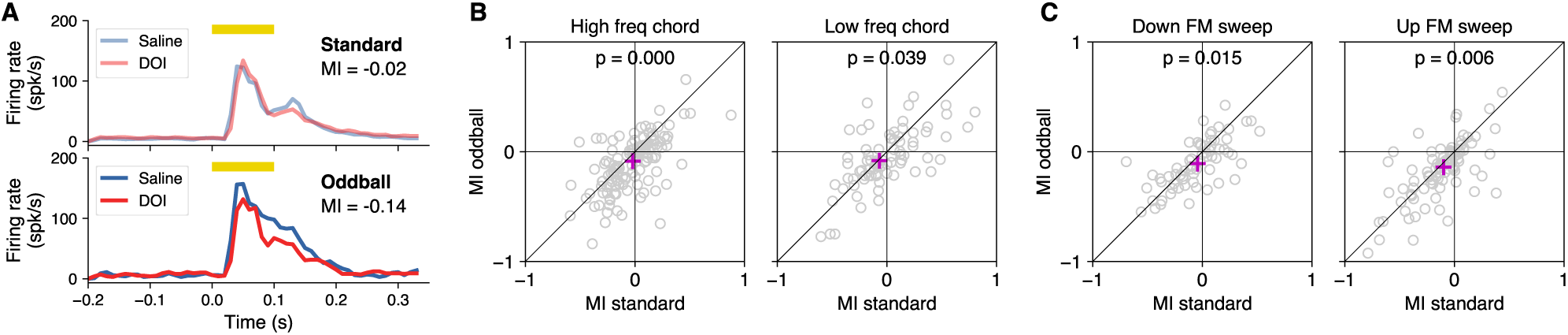
DOI reduced sensitivity to oddball sounds. **(A)** Responses of the neuron from Fig. 4F to the up-sweep FM sound when presented as the standard (top) or the oddball (bottom). Responses were almost identical between DOI and saline conditions for standards, but smaller under DOI compared to saline for oddballs. **(B)** There was a larger reduction in responses to oddballs compared to standards after DOI, for both high- and low-frequency chords. Each dot represents the modulation index (MI) from saline to DOI for each neuron. The magenta cross indicates the median value across neurons. **(C)** There was a larger reduction in responses to oddballs compared to standards after DOI, for both down- and up-sweep FM sound. Data presented as in (B).

Finally, to determine whether the reduction in responses to oddball sounds could be attributed to general response adaptation over the session, rather than the specific effects of DOI, we compared changes in responses from saline to DOI with changes from pre-injection to saline conditions. For this analysis, we used all cells that responded to the stimulus (regardless of whether their OEI was stable between pre-injection and saline). We found that there was no statistically significant change in the response to the oddball between the periods before and after saline for any stimuli, but a statistically significant change between saline and DOI for all but one stimulus tested (high-freq, *N* = 523, Pre-Saline: *p* = 0.49, Saline-DOI: *p* = 0.004; low-freq, *N* = 510, Pre-Saline: *p* = 0.58, Saline-DOI: *p <* 0.001; down-sweep, *N* = 469, Pre-Saline: *p* = 0.13, Saline-DOI: *p* = 0.16; up-sweep, *N* = 514, Pre-Saline: *p* = 0.82, Saline-DOI: *p <* 10*^−^*^6^, Wilcoxon signed-rank test). We also found that the change in oddball response between saline and DOI was more negative than the change between pre-injection and saline conditions (high-freq: MI(Sal-DOI) = *−*0.065 *vs.* MI(Pre-Sal) = *−*0.054, *p* = 0.18, low-freq: MI(Sal-DOI) = *−*0.078 *vs.* MI(Pre-Sal) = *−*0.030, *p* = 0.001, down-sweep: MI(Sal-DOI) = *−*0.072 *vs.* MI(Pre-Sal) = *−*0.034, *p* = 0.006, up-sweep: MI(Sal-DOI) = *−*0.106 *vs.* MI(Pre-Sal) = *−*0.039, *p <* 10*^−^*^6^, Wilcoxon signed-rank test). Together, these results indicate that the observed reduction in sensitivity to oddballs is primarily due to the effects of DOI and not merely the result of stimulus adaptation.

### The effects of DOI were present after repeated administration

Because our experiments included repeated administration of DOI to each animal tested, we investigated the possibility of drug tolerance, which would manifest as a smaller effect of DOI during the last experimental sessions compared to the first ones. To evaluate this possibility, we estimated the oddball enhancement index (OEI) for cells recorded during the first two sessions of each animal and compared this index to that calculated for cells recorded during the last three sessions (Fig. 6). The decision to use this specific number of sessions was driven by the goal of balancing the number of recorded cells between early and late groups.

**Figure 6:**
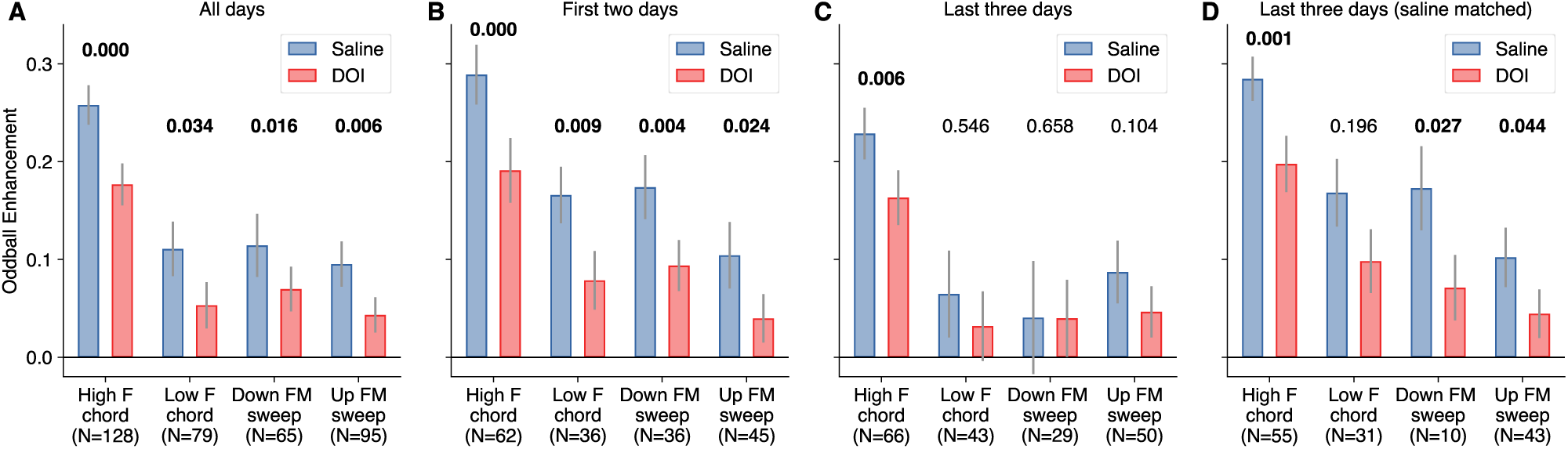
Effects of DOI were present after repeated administration. **(A)** Oddball enhancement index (OEI) under saline or DOI conditions for all neurons responsive to each sound type tested. Bars represent mean *±* standard error across neurons. Numbers above bars represent the p-value for each comparison (Wilcoxon signed-rank test). **(B)** OEI for neurons recorded during the first two sessions from each mouse. **(C)** OEI for neurons recorded during the last two sessions from each mouse. **(D)** OEI for a subset of neurons recorded during the last two sessions, such that the average OEI during saline matches that observed during the first two days.

We found that the results from the first two sessions of each mouse (Fig. 6B) were as reliable as the effects for all cells combined. In contrast, the effects observed during the last sessions appeared much smaller, reaching statistical significance for only one sound type (Fig. 6C), despite the comparable number of cells recorded between early and late sessions. However, further analysis showed that the effect of DOI (the difference in OEI between saline and DOI conditions for each neuron) during early sessions was not statistically different from the effect during late sessions (high-freq: *p* = 0.54; low-freq: *p* = 0.18; down-sweep: *p* = 0.10; up-sweep: *p* = 0.62, Mann-Whitney U rank test). Moreover, we noticed that the OEI during the saline control condition was consistently lower in cells recorded during the last three days, suggesting that the apparent differences between early and late sessions could simply reflect the recording of a sample of cells with lower OEI. To address this possibility, we created a subset of neurons from those recorded in late sessions such that this new subset would have an average OEI during saline similar to that of neurons recorded in early sessions (Fig. 6D). The results for this new subset of neurons were comparable to those observed during the first two days, except for one condition not reaching statistical significance. Moreover, a comparison of the effect of DOI between early sessions and these saline-matched sessions yielded no statistically significant difference (high-freq: *p* = 0.93; low-freq: *p* = 0.46; down-sweep: *p* = 0.57; up-sweep: *p* = 0.95, Mann-Whitney U rank test). Furthermore, performing this resampling process one hundred times, resulted in the median OEI across cells for the saline condition always being larger than the median OEI under DOI. Overall, these results suggest that the effects of DOI were present even after repeated administration of the compound, provided the cells had relatively high OEI under the control condition.

## Discussion

In this study, we evaluated changes in the activity of auditory cortical neurons when an animal is under the influence of the psychedelic 2,5-Dimethoxy-4-iodoamphetamine (DOI), a serotonergic hallucinogen. During the presentation of sounds, we found that the average firing rate of auditory cortical neurons decreased under DOI influence. This effect included a reduction in spontaneous firing, in the absolute sound-evoked firing rates, and in the relative evoked response with respect to baseline firing (Fig. 3G). We also found that in the presence of DOI, the trial-by-trial response variability increased but the frequency tuning of auditory neurons was mostly preserved across the population. When evaluating changes in sound-evoked responses during an oddball paradigm (where sequences of sounds include expected and unexpected stimuli), we found that the classically observed difference in neuronal responses between oddball and standard sounds was reduced, and that this reduction was primarily driven by a decrease in responses to oddball stimuli, suggesting a reduction in deviance detection in these neurons.

A potential limitation of working with C57BL/6J mice is that mice from this strain are known to develop progressive hearing loss, which can start as early as 10 weeks of age (31, 32). We think however that this condition has a negligible effect in the results of our study as a wide range of studies have demonstrated that young adult mice of this strain can be used for learning about the neural mechanisms of hearing, even with imperfect hearing. For example, several studies, including some from our lab, have demonstrated that young adult C57BL/6 mice can be trained to perform auditory decision-making tasks (33, 34, 35, 36), and that auditory cortical responses from mice of this strain show the expected selectivity to acoustic features (37, 38, 39). Moreover, the comparisons between the psychedelic state and controls in this study were performed within each mouse on a given day, such that any progressive loss of hearing would affect both states equally.

Changes in neural activity in the auditory system after manipulation of the serotonergic system have been observed across several brain structures along the auditory pathway (40). A suppression of neural activity after either application of serotonin or serotonin receptor agonists has been reported in the cochlear nucleus (41), inferior collicullus (42), medial geniculate nucleus of the thalamus (43) and the auditory cortex (44). While activity suppression is more commonly observed, enhancement of activity was also seen for many cells in most of these studies. This suggests that characterizing the effects of psychedelics simply as a one-directional change on overall firing rates or a decrease in responsiveness is an insufficient model for the effects of psychodelics on sound processing. Moreover, a recent study in mice found that a systemic dose of psilocybin, a serotonergic psychedelic, resulted in an initial increase in movement and neural responses in the auditory cortex followed by a decrease in responses about 30 minutes later (45), suggesting that the dynamical nature of the psychedelic effect may also need to be integrated when modeling their effects on sound processing. The timeline of our measurements matches the late stage of that study and yields a similar decrease in responses. Importantly, our study demonstrates that the effects on neural activity in the auditory cortex are not fully explained by changes in behaviors such as running, since the overall decreases in activity were observed in running as well as not-running trials.

A potential mechanism for the perceptual auditory distortions that occur with psychedelics is a consistent sharpening or broadening of frequency tuning. Our data, however, indicates that on average there is no consistent direction for changes in tuning. This result is consistent with observations that psilocybin does not significantly change pure-tone frequency selectivity of auditory cortical neurons in mice (45). Similarly, experiments evaluating the effects of systemic DOI in the visual in mice found that the orientation selectivity was preserved after DOI (16). Together, these results suggest that the mechanisms behind psychedelic-induced perceptual distortions go beyond changes in basic features of sensory processing such as stimulus tuning.

Hallucination resulting from schizophrenia and other mental disorders are hypothesized to arise from a change in the balance between external (bottom-up) and internal (top-down) signals in the brain (46). This mechanism has also been proposed as a key feature of psychedelic action (13). In the context of our experiments, a reduction in overall firing and evoked response magnitude suggests a dampening of bottom-up sensory input. This could be interpreted as a decreased ability of the auditory cortex to faithfully represent incoming auditory stimuli. However, our data also suggests alterations in top-down modulation. For instance, the reduced difference between oddball and standard responses, particularly the greater reduction in oddball responses, implies a disruption in deviance detection which relies on top-down predictive processes.

In humans, the oddball paradigm is commonly used to estimate prediction error signaling by measuring the mismatch negativity (MMN) of event-related potentials or fields, an enhancement in the negative deflection of the waveform in deviant *vs.* standard stimuli (47, 48). Manipulating the serotonergic system in humans via acute tryptophan depletion (ATD), which reduces the brain synthesis of serotonin, resulted in an increased MMN (49), consistent with a role of the serotonergic system in deviance detection as observed in our study. Moreover, a study using lysergic acid diethylamide (LSD), a serotonergic hallucinogen, found a decrease in the MMN effect after administration of this compound (50). That study found a decrease in the response to deviant stimuli under LSD, consistent with our results for DOI, although they also found an enhanced response to the standard stimuli, something not present in our data. Studies using other serotonergic hallucinogens have also found significant decreases in evoked responses in humans, but only minimal effects on the MMN when using psilocybin (51, 52, 53) or DMT (54). These various results suggest that the effects of psychedelics on deviance detection may not be the same for all compounds or may depend on specific experimental conditions.

There are at least two mechanisms that can explain the enhanced responses to oddball sounds compared to standard sounds under normal conditions: an adaptation to standard sounds and/or the neural representation of prediction errors. The first mechanism is based on the principle of neural adaptation or habituation, such that when a sound is repeatedly presented as a standard, neurons responding to that sound may decrease their firing rate over time (due to either short-term synaptic depression, activation of local inhibitory circuits, or depletion of readily releasable neurotransmitter pools). Previous studies suggest that this adaptation does not fully account for the effects observed in oddball paradigms (14). A second mechanism is that of prediction error representation, based on predictive coding theories of brain function. The brain constantly generates predictions about incoming sensory input based on recent history and context, and neurons in the auditory cortex may encode the difference between the predicted input and the actual input (*i.e.*, the prediction error). In this framework, responses to standard sounds are reduced because they are well-predicted, and oddball sounds elicit larger responses because they violate the prediction, generating a large prediction error. This mechanism involves both bottom-up sensory input and top-down predictive signals. Considering these mechanisms, we can interpret the effects of DOI on oddball responses in several ways: (1) if DOI primarily affects adaptation mechanisms, it might interfere with the normal habituation to standard sounds, reducing the contrast between standard and oddball responses; (2) if DOI impacts predictive coding processes, it could disrupt the generation or propagation of prediction errors, leading to a reduced differentiation between expected and unexpected stimuli; (3) DOI might affect both mechanisms, potentially through its action on serotonin receptors, which are known to modulate both local circuit dynamics and long-range connections involved in predictive processing.

In the context of our experiments, if DOI primarily affected adaptation, we would expect a reduction in the difference between oddball and standard responses, as observed in our data. However, we might expect this to occur primarily through an increase in the response to standards (due to reduced adaptation) rather than a decrease in the response to oddballs. The overall reduction in responses and the greater effect on oddballs suggest that something more complex is occurring. If DOI primarily affected prediction error signaling, we would expect a reduction in the difference between oddball and standard responses, but also a more pronounced effect on oddball responses, as these represent larger prediction errors. This aligns with our observation of a greater reduction in oddball responses. Moreover, the overall reduction in responsiveness is consistent with a general dampening of prediction error signaling. Therefore, we conclude that our results align more closely to the prediction error mechanisms being affected by DOI. This of course does not rule out effects on adaptation, but rather, it suggests that the impact on prediction error signaling may be more prominent in our experimental paradigm.

## Data availability

Source data for this study are openly available at https://doi.org/10.5281/zenodo.14285875.

## Acknowledgments

We thank the Terrestrial Animal Care Services at the University of Oregon for providing excellent animal husbandry and veterinary care, and George Vengrovski for assistance in the development of implanted devices. Generative AI tools, such as GPT-4 and Claude 3, were used for revising the text of the manuscript. These tools were used in a manner that does not conflict with APS ethical policies and the authors take full responsibility for the content.

## Author contributions

SJ conceived the project. MH and JLM conducted the experiments. MH and SJ analyzed the data. SJ wrote the paper.

## References

[1] Alexander T. Shulgin and Ann Shulgin. Pihkal: A Chemical Love Story. Transform Press, Berkeley, CA, 1991.

[2] Alexander T. Shulgin and Ann Shulgin. Tihkal: The Continuation. Transform Press, Berkeley, CA, 1997.

[3] Collin M. Reiff, Elon E. Richman, Charles B. Nemeroff, Linda L. Carpenter, Alik S. Widge, Carolyn I. Rodriguez, Ned H. Kalin, William M. McDonald, and and the Work Group on Biomarkers and Novel Treatments, a Division of the American Psychiatric Association Council of Research. Psychedelics and Psychedelic-Assisted Psychotherapy. American Journal of Psychiatry, 177(5):391–410, May 2020.

[4] Jennifer M. Mitchell and Brian T. Anderson. Psychedelic therapies reconsidered: Compounds, clinical indications, and cautious optimism. Neuropsychopharmacology, 49(1):96–103, January 2024.

[5] David E. Nichols. Psychedelics. Pharmacological Reviews, 68(2):264–355, April 2016.

[6] Katrin H. Preller and Franz X. Vollenweider. Phenomenology, Structure, and Dynamic of Psychedelic States. In Adam L. Halberstadt, Franz X. Vollenweider, and David E. Nichols, editors, Behavioral Neurobiology of Psychedelic Drugs, pages 221–256. Springer, Berlin, Heidelberg, 2018.

[7] Jacob S. Aday, Julia R. Wood, Emily K. Bloesch, and Christopher C. Davoli. Psychedelic drugs and perception: A narrative review of the first era of research. Reviews in the Neurosciences, 32(5):559–571, July 2021.

[8] Marc Wittmann, Olivia Carter, Felix Hasler, B. Rael Cahn, Ulrike Grimberg, Philipp Spring, Daniel Hell, Hans Flohr, and Franz X. Vollenweider. Effects of psilocybin on time perception and temporal control of behaviour in humans. Journal of Psychopharmacology, 21(1):50–64, January 2007.

[9] Alex C. Kwan, David E. Olson, Katrin H. Preller, and Bryan L. Roth. The neural basis of psychedelic action. Nature Neuroscience, 25(11):1407–1419, November 2022.

[10] Franz X Vollenweider and Mark A Geyer. A systems model of altered consciousness: Integrating natural and drug-induced psychoses. Brain Research Bulletin, 56(5):495–507, November 2001.

[11] Robin L. Carhart-Harris, Robert Leech, Peter J. Hellyer, Murray Shanahan, Amanda Feilding, Enzo Tagliazucchi, Dante R. Chialvo, and David Nutt. The entropic brain: A theory of conscious states informed by neuroimaging research with psychedelic drugs. Frontiers in Human Neuroscience, 8, February 2014.

[12] Katrin H. Preller, Adeel Razi, Peter Zeidman, Philipp Stämpfli, Karl J. Friston, and Franz X. Vollenweider. Effective connectivity changes in LSD-induced altered states of consciousness in humans. Proceedings of the National Academy of Sciences, 116(7):2743–2748, February 2019.

[13] R. L. Carhart-Harris and K. J. Friston. REBUS and the Anarchic Brain: Toward a Unified Model of the Brain Action of Psychedelics. Pharmacological Reviews, 71(3):316–344, July 2019.

[14] Israel Nelken. Stimulus-specific adaptation and deviance detection in the auditory system: Experiments and models. Biological Cybernetics, 108(5):655–663, October 2014.

[15] Rik J. J. van Daal, C. aăatay Aydin, Frédéric Michon, Arno A. A. Aarts, Michael Kraft, Fabian Kloosterman, and Sebastian Haesler. Implantation of Neuropixels probes for chronic recording of neuronal activity in freely behaving mice and rats. Nature Protocols, 16(7):3322–3347, July 2021.

[16] Angie M. Michaiel, Philip R. L. Parker, and Cristopher M. Niell. A Hallucinogenic Serotonin-2A Receptor Agonist Reduces Visual Response Gain and Alters Temporal Dynamics in Mouse V1. Cell Reports, 26(13):3475–3483.e4, March 2019.

[17] Jonathan C. Gewirtz and Gerard J. Marek. Behavioral Evidence for Interactions between a Hallucinogenic Drug and Group II Metabotropic Glutamate Receptors. Neuropsychopharmacology, 23(5):569–576, November 2000.

[18] Clint E. Canal and Drake Morgan. Head-twitch response in rodents induced by the hallucinogen 2,5-dimethoxy-4-iodoamphetamine: A comprehensive history, a re-evaluation of mechanisms, and its utility as a model. Drug testing and analysis, 4(7-8):556–576, 2012.

[19] Abdullah Al Shoyaib, Sabrina Rahman Archie, and Vardan T. Karamyan. Intraperitoneal Route of Drug Administration: Should it Be Used in Experimental Animal Studies? Pharmaceutical Research, 37(1):12, December 2019.

[20] Quanxin Wang, Song-Lin Ding, Yang Li, Josh Royall, David Feng, Phil Lesnar, Nile Graddis, Maitham Naeemi, Benjamin Facer, Anh Ho, Tim Dolbeare, Brandon Blanchard, Nick Dee, Wayne Wakeman, Karla E. Hirokawa, Aaron Szafer, Susan M. Sunkin, Seung Wook Oh, Amy Bernard, John W. Phillips, Michael Hawrylycz, Christof Koch, Hongkui Zeng, Julie A. Harris, and Lydia Ng. The Allen Mouse Brain Common Coordinate Framework: A 3D Reference Atlas. Cell, 181(4):936–953.e20, May 2020.

[21] Atika Syeda, Lin Zhong, Renee Tung, Will Long, Marius Pachitariu, and Carsen Stringer. Facemap: A framework for modeling neural activity based on orofacial tracking. Nature Neuroscience, 27(1):187–195, January 2024.

[22] Marius Pachitariu, Shashwat Sridhar, Jacob Pennington, and Carsen Stringer. Spike sorting with Kilosort4. Nature Methods, 21(5):914–921, May 2024.

[23] Cyrille Rossant, Shabnam N. Kadir, Dan F. M. Goodman, John Schulman, Maximilian L. D. Hunter, Aman B. Saleem, Andres Grosmark, Mariano Belluscio, George H. Denfield, Alexander S. Ecker, Andreas S. Tolias, Samuel Solomon, György Buzsáki, Matteo Carandini, and Kenneth D. Harris. Spike sorting for large, dense electrode arrays. Nature Neuroscience, 19(4):634–641, April 2016.

[24] Adam L. Halberstadt and Mark A. Geyer. Characterization of the head-twitch response induced by hallucinogens in mice: Detection of the behavior based on the dynamics of head movement. Psychopharmacology, 227(4):10.1007/s00213-013-3006-z, June 2013.

[25] Adam L. Halberstadt, Iris van der Heijden, Michael A. Ruderman, Victoria B. Risbrough, Jay A. Gingrich, Mark A. Geyer, and Susan B. Powell. 5-HT2A and 5-HT2C Receptors Exert Opposing Effects on Locomotor Activity in Mice. Neuropsychopharmacology, 34(8):1958–1967, July 2009.

[26] David M. Schneider, Anders Nelson, and Richard Mooney. A synaptic and circuit basis for corollary discharge in the auditory cortex. Nature, 513(7517):189–194, September 2014.

[27] Nachum Ulanovsky, Liora Las, and Israel Nelken. Processing of low-probability sounds by cortical neurons. Nature Neuroscience, 6(4):391–398, April 2003.

[28] Johannes J. Letzkus, Steffen B. E. Wolff, Elisabeth M. M. Meyer, Philip Tovote, Julien Courtin, Cyril Herry, and Andreas Lüthi. A disinhibitory microcircuit for associative fear learning in the auditory cortex. Nature, 480(7377):331–335, December 2011.

[29] Tyler L. Gimenez, Maja Lorenc, and Santiago Jaramillo. Adaptive categorization of sound frequency does not require the auditory cortex in rats. Journal of Neurophysiology, 114(2):1137–1145, August 2015.

[30] Sebastian Ceballo, Zuzanna Piwkowska, Jacques Bourg, Aurélie Daret, and Brice Bathellier. Targeted Cortical Manipulation of Auditory Perception. Neuron, 104(6):1168–1179.e5, December 2019.

[31] Elizabeth M. Keithley, Cecilia Canto, Qing Yin Zheng, Nathan Fischel-Ghodsian, and Kenneth R. Johnson. Age-related hearing loss and the *ahl* locus in mice. Hearing Research, 188(1):21–28, February 2004.

[32] James R. Ison, Paul D. Allen, and William E. O’Neill. Age-Related Hearing Loss in C57BL/6J Mice has both Frequency-Specific and Non-Frequency-Specific Components that Produce a Hyperacusis-Like Exaggeration of the Acoustic Startle Reflex. Journal of the Association for Research in Otolaryngology, 8(4):539–550, December 2007.

[33] Santiago Jaramillo and Anthony M. Zador. Mice and rats achieve similar levels of performance in an adaptive decision-making task. Frontiers in Systems Neuroscience, 8, September 2014.

[34] Lan Guo, William I. Walker, Nicholas D. Ponvert, Phoebe L. Penix, and Santiago Jaramillo. Stable representation of sounds in the posterior striatum during flexible auditory decisions. Nature Communications, 9(1):1534, April 2018.

[35] Lan Guo, Jardon T. Weems, William I. Walker, Anastasia Levichev, and Santiago Jaramillo. Choice-Selective Neurons in the Auditory Cortex and in Its Striatal Target Encode Reward Expectation. Journal of Neuroscience, 39(19):3687–3697, May 2019.

[36] Anna A. Lakunina, Nadav Menashe, and Santiago Jaramillo. Contributions of Distinct Auditory Cortical Inhibitory Neuron Types to the Detection of Sounds in Background Noise. eNeuro, 9(2), March 2022.

[37] Nicholas D. Ponvert and Santiago Jaramillo. Auditory Thalamostriatal and Corticostriatal Pathways Convey Complementary Information about Sound Features. Journal of Neuroscience, 39(2):271–280, January 2019.

[38] Anna A. Lakunina, Matthew B. Nardoci, Yashar Ahmadian, and Santiago Jaramillo. Somatostatin-Expressing Interneurons in the Auditory Cortex Mediate Sustained Suppression by Spectral Surround. Journal of Neuroscience, 40(18):3564–3575, April 2020.

[39] Jennifer L. Mohn, Melissa M. Baese-Berk, and Santiago Jaramillo. Selectivity to acoustic features of human speech in the auditory cortex of the mouse. Hearing Research, 441:108920, January 2024.

[40] L. M. Hurley and I. C. Hall. Context-dependent modulation of auditory processing by serotonin. Hearing Research, 279(1):74–84, September 2011.

[41] Ulrich Ebert and Joachim Ostwald. Serotonin modulates auditory information processing in the cochlear nucleus of the rat. Neuroscience Letters, 145(1):51–54, September 1992.

[42] Laura M. Hurley and George D. Pollak. Serotonin Differentially Modulates Responses to Tones and Frequency-Modulated Sweeps in the Inferior Colliculus. Journal of Neuroscience, 19(18):8071–8082, September 1999.

[43] James E. Monckton and David A. McCormick. Neuromodulatory Role of Serotonin in the Ferret Thalamus. Journal of Neurophysiology, 87(4):2124–2136, April 2002.

[44] Weiqing Ji and Nobuo Suga. Serotonergic Modulation of Plasticity of the Auditory Cortex Elicited by Fear Conditioning. Journal of Neuroscience, 27(18):4910–4918, May 2007.

[45] Adam T. Brockett and Nikolas A. Francis. Psilocybin decreases neural responsiveness and increases functional connectivity while preserving pure-tone frequency selectivity in mouse auditory cortex. Journal of Neurophysiology, 132(1):45–53, July 2024.

[46] Stephen Grossberg. How hallucinations may arise from brain mechanisms of learning, attention, and volition. Journal of the International Neuropsychological Society, 6(5):583–592, July 2000.

[47] R. Näätänen, M. Simpson, and N. E. Loveless. Stimulus deviance and evoked potentials. Biological Psychology, 14(1):53–98, February 1982.

[48] Marta I. Garrido, James M. Kilner, Klaas E. Stephan, and Karl J. Friston. The mismatch negativity: A review of underlying mechanisms. Clinical Neurophysiology, 120(3):453–463, March 2009.

[49] Seppo Kähkönen, Ville Mäkinen, Iiro P. Jääskeläinen, Sirpa Pennanen, Jyrki Liesivuori, and Jyrki Ahveninen. Serotonergic modulation of mismatch negativity. Psychiatry Research: Neuroimaging, 138(1):61–74, January 2005.

[50] Christopher Timmermann, Meg J. Spriggs, Mendel Kaelen, Robert Leech, David J. Nutt, Rosalyn J. Moran, Robin L. Carhart-Harris, and Suresh D. Muthukumaraswamy. LSD modulates effective connectivity and neural adaptation mechanisms in an auditory oddball paradigm. Neuropharmacology, 142:251–262, November 2018.

[51] Daniel Umbricht, Franz X. Vollenweider, Liselotte Schmid, Claudia Grübel, Anja Skrabo, Theo Huber, and Rene Koller. Effects of the 5-HT2A Agonist Psilocybin on Mismatch Negativity Generation and AX-Continuous Performance Task: Implications for the Neuropharmacology of Cognitive Deficits in Schizophrenia. Neuropsychopharmacology, 28(1):170–181, January 2003.

[52] André Schmidt, Rosilla Bachmann, Michael Kometer, Philipp A. Csomor, Klaas E. Stephan, Erich Seifritz, and Franz X. Vollenweider. Mismatch Negativity Encoding of Prediction Errors Predicts S-ketamine-Induced Cognitive Impairments. Neuropsychopharmacology, 37(4):865–875, March 2012.

[53] Anna Bravermanová, Michaela Viktorinová, Filip Tylš, Tomáš Novák, Renáta Androvičová, Jakub Korčák, Jiří Horáček, Marie Balíková, Inga Griškova-Bulanova, Dominika Danielová, Přemysl Vlček, Pavel Mohr, Martin Brunovský, Vlastimil Koudelka, and Tomáš Páleníček. Psilocybin disrupts sensory and higher order cognitive processing but not pre-attentive cognitive processing—study on P300 and mismatch negativity in healthy volunteers. Psychopharmacology, 235(2):491–503, February 2018.

[54] Karsten Heekeren, Jörg Daumann, Anna Neukirch, Carsten Stock, Wolfram Kawohl, Christine Norra, Till D. Waberski, and Euphrosyne Gouzoulis-Mayfrank. Mismatch negativity generation in the human 5HT2A agonist and NMDA antagonist model of psychosis. Psychopharmacology, 199(1):77–88, July 2008.

